# Genetic association between posterior parietal cortex and intelligence

**DOI:** 10.1101/2025.03.21.644693

**Authors:** Dong Yang, Xushen Yang, Jikan Peng, Junfeng Lu, Xiaotao Hao, Shubin Guo, Jiaojiao Li, Runze Chen, Chenshi Xu, Lijun Xiang, Tian Xu

## Abstract

The identification of intelligence-associated loci raises questions for functional association of brain regions and for different intelligence concepts of either a general capability or discrete abilities. By generating single-nucleus transcriptomic atlas of mouse posterior parietal cortex (PPC), which is implicated in quantification behavior, we find that human intelligence-associated gene homologs are significantly enriched in certain specific neurons within the PPC. We also find 73% of the human intelligence gene homologs are highly enriched in PPC. In addition to PPC, several regions at different locations, not cortex next to PPC, are also significantly associated with intelligence genes. The separate locations of these cortex regions and their regional-specific expressed intelligence genes raise the possibility for discrete intelligence capabilities. Animal behavior analysis reveals that mutations for two top intelligence-associated genes with specific expression in either PPC or entorhinal cortex (ENT) affect quantification or learning and memory, respectively. Finally, five patients with PPC resection in the left hemisphere also display quantification defects, but not other cognitive abilities. Together, our studies provide the first genetic and functional evidence for the involvement of PPC in intelligence. Our data also support the model of multiple intelligence over a general capability for intelligence.

## INTRODUCTION

Recent large-scale human genome-wide studies have significantly advanced our knowledge of the genetic basis of intelligence^1–3^. However, the identified loci contain hundreds of functionally distinct genes that do not immediately suggest brain regions and cell types for further experiments. The brain comprises diverse regions, which are further composed of a variety of cell types^4,5^. It would be important to map the intelligence-associated genes to specific brain regions and cell types.

Intelligence is often considered as a general capability^7,8^ and is commonly inferred by a holistic intelligence test including the assessment of quantification ability^6^. On the other hand, Howard Gardner and others proposed the multiple intelligences theory, stating that intelligence is not a general capability, but rather a set of independent abilities such as logic and quantification abilities^9^.

Integrating the human genomic findings, transcriptomic brain cell taxonomy and animal ethological analysis will provide deeper insights. We have therefore systematically generated a single-cell gene expression atlas of mouse PPC, a region implicated in quantification ability. We found that a subset of the intelligence-associated genes is consistently mapped to PPC neurons, while the others are mapped to several separate cortical regions. Analysis of mouse mutants for two intelligence-associated genes and patients with PPC resection provides further evidence for the association between PPC and intelligence and argues for intelligence of multiple separable abilities.

## RESULTS

### Atlas of cell-type across the PPC using sn RNA-seq

To generate a high-resolution single-cell atlas of PPC, 10× Genomics Chromium droplet-based single-nucleus RNA-seq was used to profile three library replicates, each derived from cells collected from 25 adult mice (Figure 1A, Methods). A total of 39,843 high-quality single-cell transcriptomes are obtained. We perform unsupervised graph-based clustering and identify 24 cell populations in PPC, each with a distinct gene expression pattern (Figures 1B-D, Figure S1). Based on their significantly enriched genes, we have annotated the glutamatergic excitatory neurons (*Slc17a7*) and the GABAergic inhibitory neurons (*Gad2*). In addition, non-neuronal cells include oligodendrocytes (*Mog*), oligodendrocyte precursor cells (*Vcan*), astrocytes (*Gja1*), microglial cells (*C1qa*) and endothelial cells (*Vtn*) (Figures 1C-D, Figure S1). The proportion of glutamatergic neurons (69%) far exceeds that of the GABAergic neurons (10%) (Figure 1E). The relative cell numbers of these classes are consistent across biological replicates (Figures 1E-G), indicating that the proportions and diversity of cells are similarly regulated during development to maintain the function of PPC.

**Figure 1.**
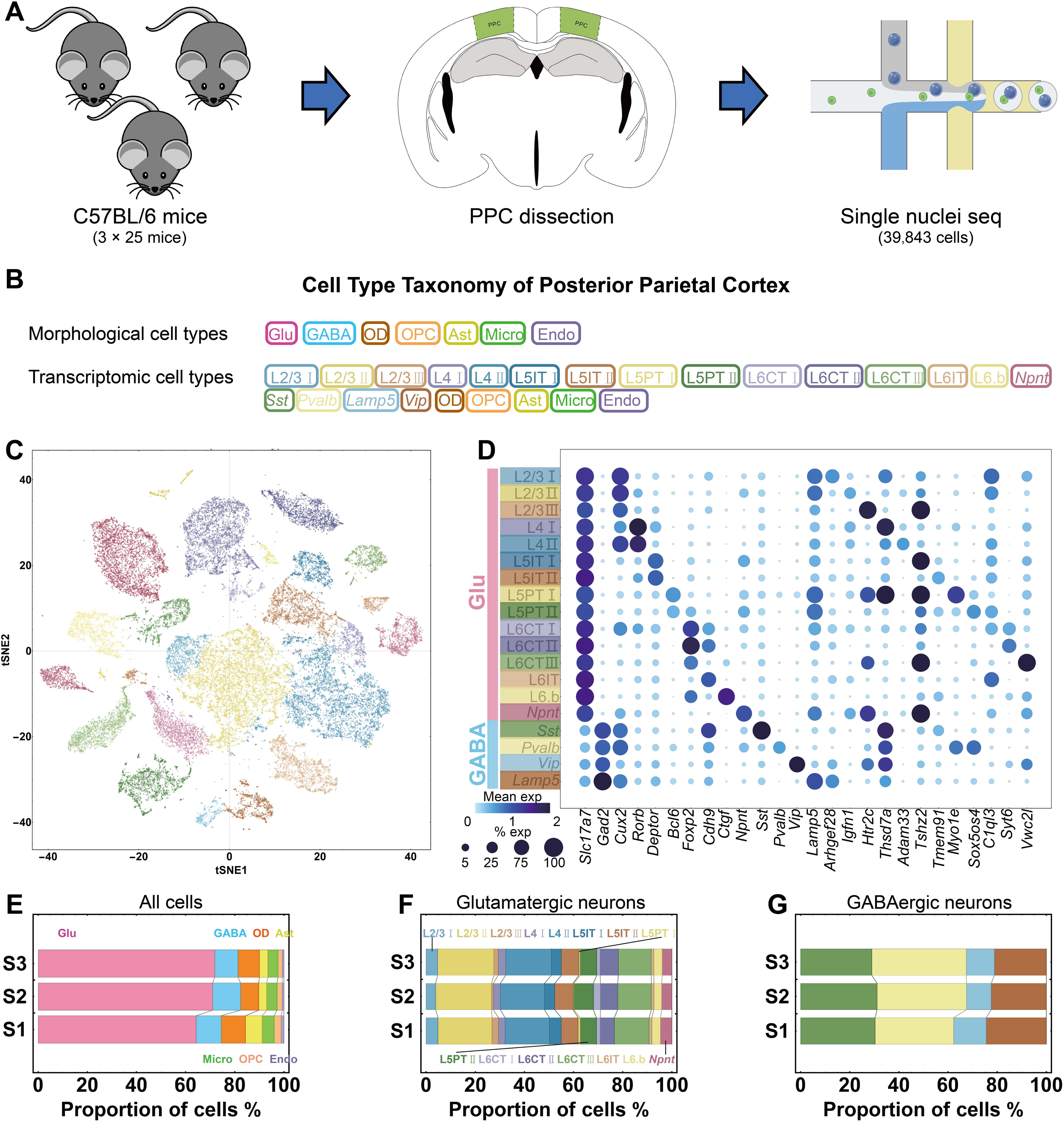
Profiling of cell types in PPC by snRNA-seq. (**A**) Experimental workflow of snRNA-seq profiling of PPC. (**B**) Morphological and transcriptomic cell type taxonomy of PPC. (**C**) t-SNE plot visualization of the PPC cell types. Dots represent cells, and distances between dots reflect transcriptomic similarity. Cells are colored by transcriptomic types. (**D**) Bubble heatmap showing the expression levels of the selected marker genes in PPC neurons. (**E-G**) Proportions of cells profiled according to (E) all cells, (F) Glutamatergic neurons or (G) GABAergic neurons. Abbreviations: Glu: Glutamatergic neurons; GABA: GABAergic neurons; OD: Oligodendrocytes; Ast: Astrocyte; Micro: Microglia; OPC: Oligodendrocyte precursor cells; Endo: Endothelial cell. L2/3, 4, 5, and 6: layer 2/3, 4, 5, and 6, respectively. S1-S3: 3 biological replicates.

In addition to 4 classes of GABAergic neurons and 14 classes of glutamatergic neurons that have been previously reported^5,10^, we have identified a new class of glutamatergic neurons, which specifically express *Npnt* (*Nephronectin*) (Figure 1D). Single-molecule fluorescence in situ hybridization reveals that the *Npnt* glutamatergic neurons localize in layer 2/3 and layer 5 (Figure S2, Methods). Furthermore, they are different from previously identified neurons in these layers as they express neither *Cux2* nor *Deptor* (Figure S2). Together, our data serve as a reference for transcriptomic taxonomy of cells in PPC (Figure 1).

### Genetic identification of the association between PPC and intelligence

To identify the physiological or disease processes that are associated with PPC, we apply our data to evaluate whether the genomic loci identified in the large human studies map to cell types in PPC (Figures 2A and B, Methods). We first analyze data with neurological diseases including schizophrenia^11^, attention deficit hyperactivity disorder (ADHD)^12^, Alzheimer^13^, bipolar disorder^14^, depression^15^, obsessive-compulsive disorder (OCD)^16^, and Parkinson’s disease^17^. While associations between these diseases with neuron types in other brain regions had been reported^18,19^, we do not find any enrichment signal for the cell types in PPC (Figure 2B).

**Figure 2.**
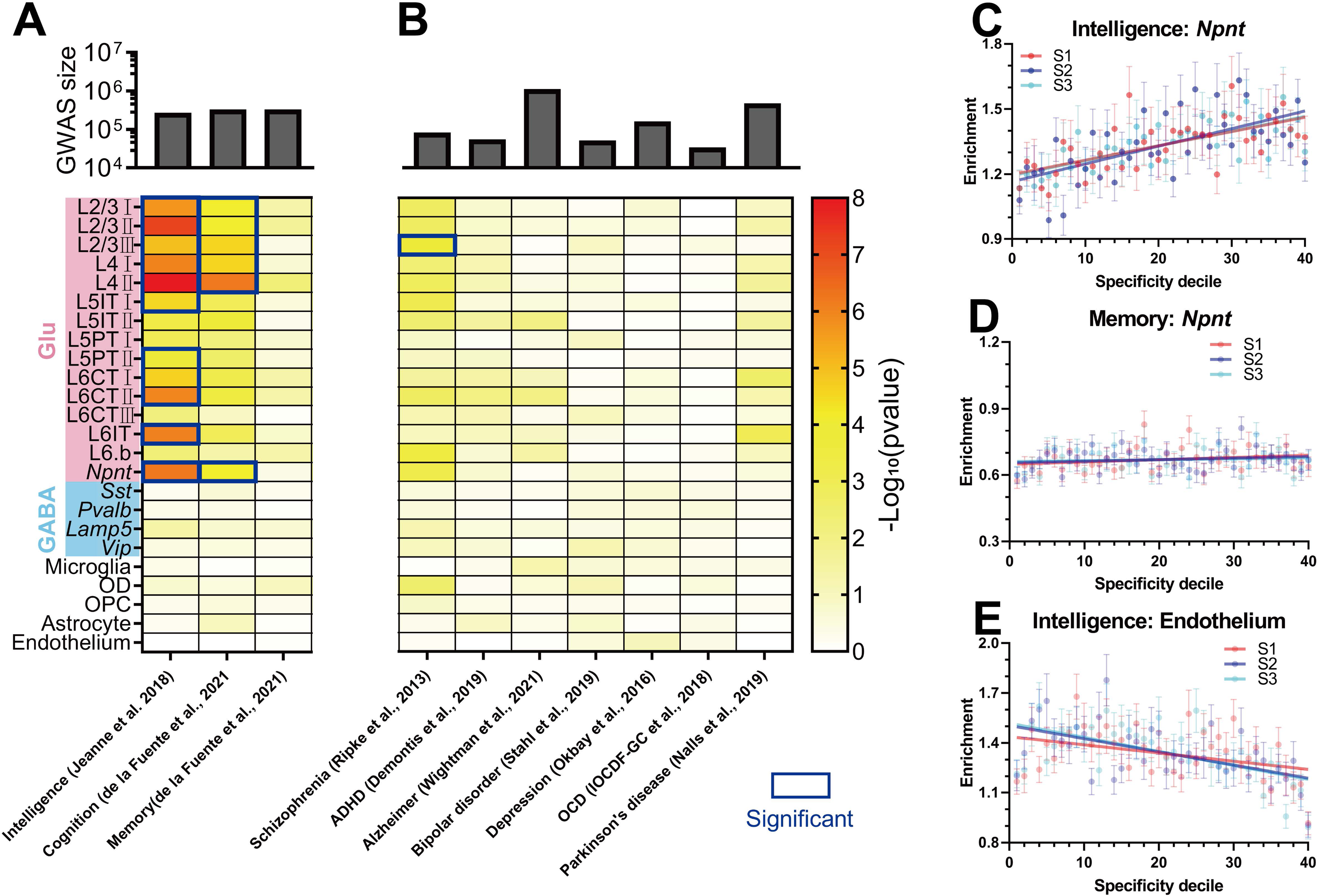
Genetic association between PPC and intelligence. (**A-B**) Heatmap showing the association *P* values of diverse human cognitive traits (A) or neurological diseases (B) with all the cell types in mouse PPC. (**C-E**) Scatterplots showing examples of the enrichment of a given trait-related genes heritability in each of the specificity deciles for a given cell types (C, positive correlation; D and E, no correlation). Lines showing the linear regression according to 3 biological repeats. S1-S3: 3 biological replicates. The enrichment (y axis) represents the mean GWAS intelligence-associated p value (-log_10_ transformed) of the genes.

To identify the neurophysiological processes that are associated with the cells in PPC, we use the largest available human GWAS data, including intelligence^20^, cognition^21^, and memory^21^. No enrichment signals are detected for the memory-associated genes in PPC (Figure 2A). While no enrichment signal is detected for the majority of the cognition-associated genes, enrichment signal is identified for a small number of the cognition-associated genes with several PPC cell types (Figure 2A). This might be due to the genes shared by cognition and intelligence. Instead, we found that the intelligence-associated genes consistently mapped to the excitatory neurons in PPC (p=6.46×10^-9^ to 5.25×10^-5^, Figure 2A). For example, we plot the enrichment of heritability for intelligence^22^ in the cell-type specificity deciles for *Npnt* glutamatergic neurons in PPC and find a significant positive relationship (p=3.69×10^-7^, Figures 2A, 2C). No such relationship is detected between *Npnt* neurons and memory (Figure 2D) or between endothelium cell and intelligence (Figure 2E).

To confirm the association between PPC and intelligence, we extend our analysis to the cortical regions adjacent to PPC, including primary visual (VISp), medial visual (VISm), and primary somatosensory (SSp)^5^. We do not detect any significant association between intelligence and these three cortical regions (Figure 3A). Together, these data highlight the significance of PPC in its association with intelligence.

**Figure 3.**
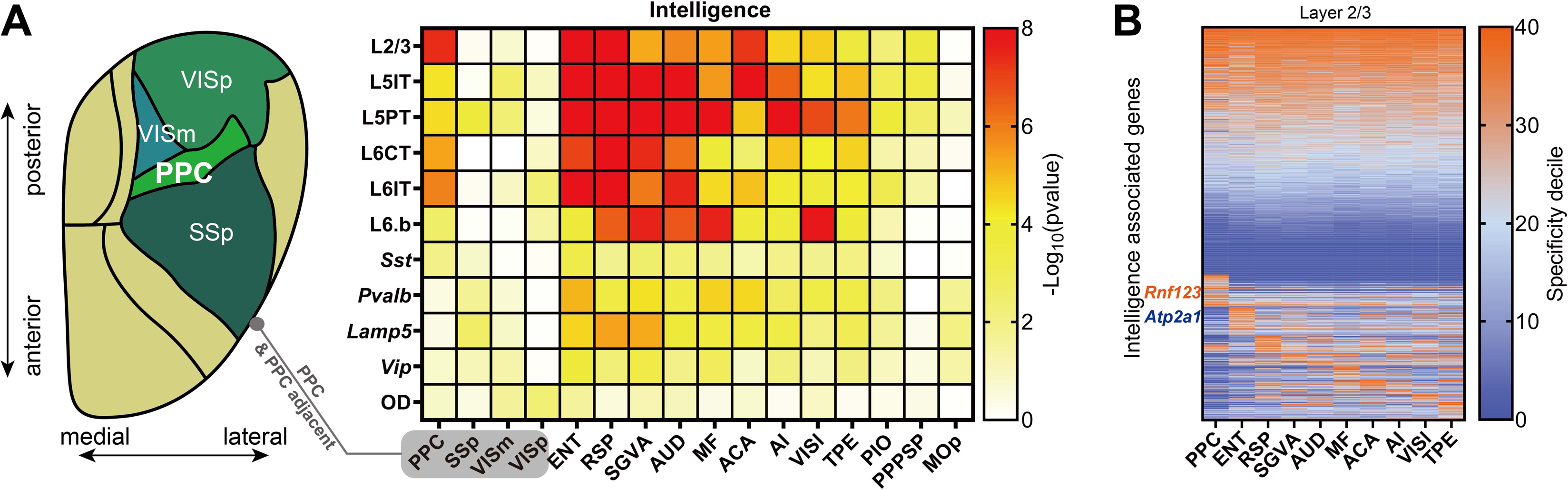
Genetic association between intelligence and additional cortical regions. **(A)** Heatmap (right) of the association between intelligence and multiple cortical regions. 2D flatmap (left) representation of cortical regions adjacent to PPC according to their position. (**B**) Heatmap of the expression patterns of intelligence genes across the intelligence-associated cortical regions in layer 2/3 neurons.

#### Genetic association between intelligence and additional cortical regions

We expand our analysis of genetic association with intelligence to additional cortical regions using single-cell atlas data^5^. In addition to PPC, several other cortical regions or combined regions are identified to have significant association with intelligence including entorhinal (ENT), retrosplenial (RSP), supplemental somatosensory-gustatory-visceral-posterior agranular insular (SGVA), auditory (AUD), secondary motor-frontal pole (MF), anterior cingulate (ACA), agranular insular (AI), lateral visual areas (VISl), and temporal association-perirhinal-ectorhinal joint regions (TPE) (Figure 3A). On the other hand, there are cortical regions do not have significant association with intelligence including primary motor cortex (MOp), parasubiculum-postsubiculum-presubiculum-subiculum-prosubiculum joint areas (PPPSP), and prelimbic-infralimbic-orbital joint areas (PIO) (Figure 3A).

### Partitioning genetic contributions for different intelligence abilities

Given multiple cortical regions are enriched with intelligence genes, we are interested in learning whether there is a subset of the intelligence genes which are specifically associated with a given cortical region. Indeed, about 36.9% of the intelligence-associated genes are expressed in a cortical region-specific manner (Figure 3B, Methods). For example, 73.5% (699/951) of the genes are expressed in PPC neurons, while 8.2% (78/951) of the intelligence-associated genes are expressed specifically in PPC (Figure 3B). Similarly, about 7.4% (70/951) of the intelligence-associated genes show specificity in ENT, an area mainly involved in learning and memory^23,24^ (Figure 3B). These observations raise the possibility that these cortical region-specific expressed subsets of intelligence genes could contribute differently to intelligence.

To functionally test this hypothesis, we therefore focus on a candidate gene with the highest association value for intelligence (*Rnf123*, p = 1.63×10^−33^)^20^ and high PPC specificity (Figures 3B, 4A). We found that mice have the ability to distinguish the amount of food (Methods, Jikan Peng et al., 2024, under submission). Significantly, *Rnf123* homozygous mutant mice completely this ability (p = 0.002) (Figure 4B). Interestingly, further experiments show that the *Rnf123* mutant mice display normal learning and memory capacity (Figure 4C, Methods). To further explore this hypothesis, we have examined another top intelligence-associated gene (p = 4.62×10^-23^), *Atp2a1* (Figure 3B, 4A), which is enriched in ENT, but not in PPC. Correspondingly, while *Atp2a1* homozygous mutants are lethal^25^, the *Atp2a1* heterozygous mice exhibit a significant deficit in learning and memory ability (p = 0.0002), but normal in quantification of food (Figure 4B, C). Analysis of the locomotion speed and behaviors in y maze confirm the normal motor function and working memory of these two mouse strains (Figures S3A, B). These experiments not only provided functional genetic data confirming the involvement of PPC in quantification behavior, but also supported the hypothesis that the cortex-region-specific expressed intelligence genes are involved in distinctive functions.

**Figure 4.**
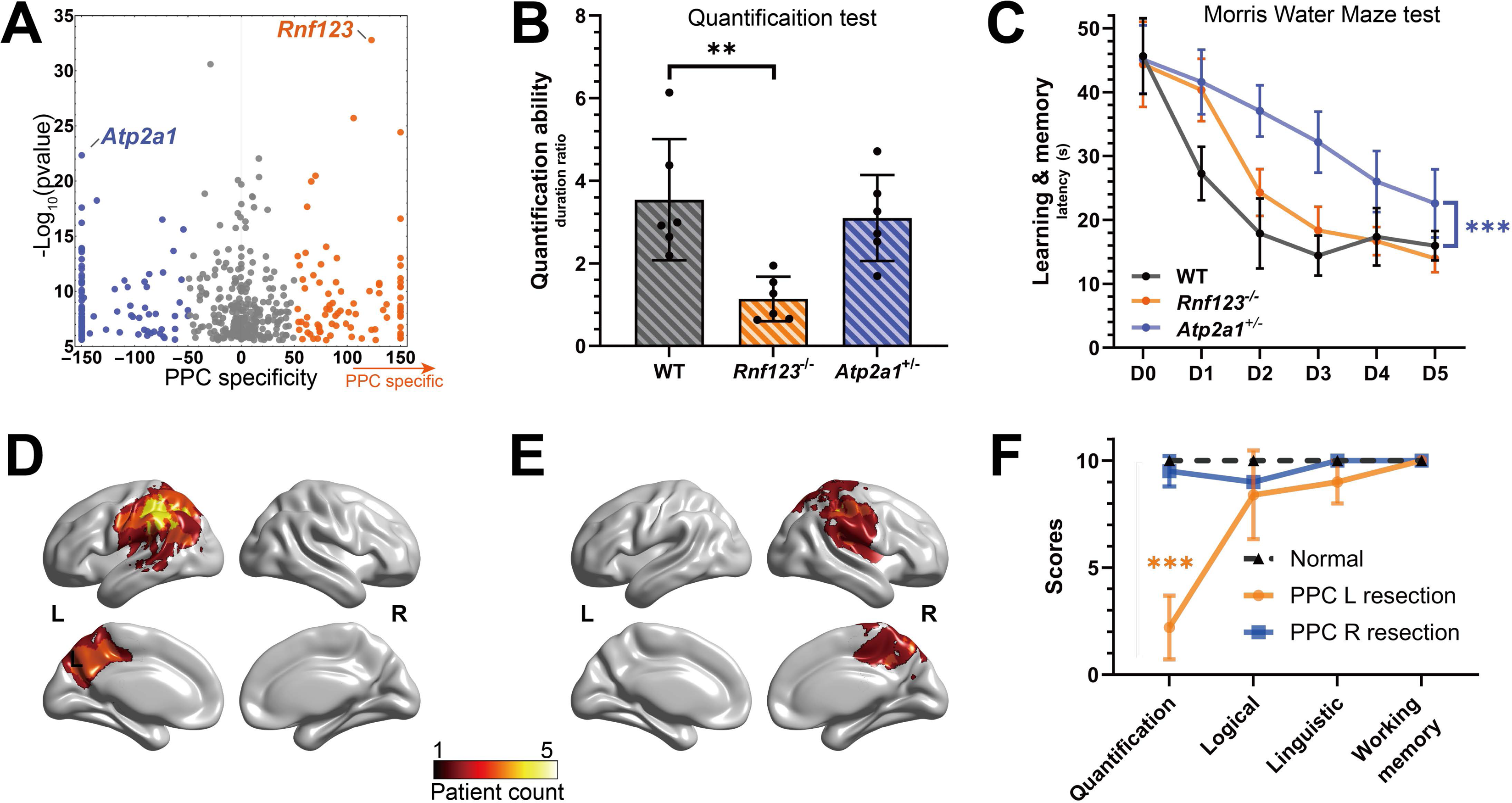
Partitioning genetic contributions for quantification intelligence. (**A**) Dot plot quantifying the specificity of intelligence genes in PPC compared to other intelligence associated cortical regions. *Rnf123* and *Atp2a1* are selected for animal study due to their high association with intelligence (y axis) and their selective expression patterns (x axis). (**B, C**) Mouse behavior tests showing the deficit of *Rnf123^-/-^* mice in quantification ability (B) and abnormality of *Atp2a1^+/-^* mice in learning memory ability (C). Quantification ability is calculated by the ratio of the duration time in the area with 80g and 20g food. Mann-Whitney U test. (**D-F**) Patients with PPC lesions and quantification ability tests. Overlay maps of lesions including PPC in the left hemisphere in patients 1-5 (D) and in the right hemisphere in patients 6-7 (E). The color scale represents the number of patients with a lesion in a particular voxel. (F) Test scores from multiple human abilities showing the significant defect of quantification ability in the patients with lesions involving PPC in the left hemisphere. The scores represent the number of correct answers out of 10 questions.

We therefore investigate the involvement of PPC in the quantification ability of mice. We find chemogenetic inhibition of PPC neurons eliminate the quantification preference (Jikan Peng et al., 2024, under submission). We further analyzed a group of 7 patients with brain resection lesions involving in PPC (Figures 4D, E, Figure S4, Methods). Five of the patients had lesions in the left hemisphere, and the only common lesion region in these patients is PPC (Figure 4D). Interestingly, all these five patients with PPC lesions displayed the deficient in quantification ability, but normal in logical, linguistic, and learning memory functions (Figure 4F). No quantification defect was detected in the two patients with lesions associated with PPC in the right hemisphere (Figure 4F). Together, both mouse and human data support the involvement of PPC in quantification ability but not in other cognitive abilities and argue for partitioning intelligence into multiple abilities.

## DISCUSSION

Recent genomic studies have significantly advanced our understanding of genetic contribution to intelligence by the identification of a collection of intelligence-associated genes^1–3^, revealing how these genes are expressed in the brain could help to study how they contribute to physiological functions and behaviors. Our analysis here presents a comprehensive single cell transcriptomic atlas of mouse PPC and define 24 cell types with their specific transcriptomic patterns and spatial arrangement. This landscape of PPC serves as a useful reference to explore neural biology. Linking our data to genomic findings for human cognitive processes or psychiatric disorders, we have identified significant enrichment of intelligence-associated genes in the PPC neurons, but not with genes for other cognitive processes or psychiatric disorders. Furthermore, such genetic association between intelligence and PPC is not presented for the other cortical regions adjacent to PPC. Finally, alteration of the top intelligence-associated gene, *Rnf123*, which specifically expresses in PPC, disrupts quantification behavior in mice. Consistent with the animal results, five patients with PPC resection lesions in the left hemisphere displays quantification defects. This work has provided, for the first time, the genetic and functional data associating PPC with quantification intelligence.

While intelligence is commonly measured by tests composed with different cognitive skills or behaviors^6^, it was commonly thought as a general ability which influences the performance of all cognitive abilities for more than a centry^7^. However, Gardner and others had presented a dissenting view since 1983 arguing for intelligence as a set of independent abilities such as quantification, linguistic, and logic capabilities^9^. We have comprehensively mapped intelligence-associated genes to the entire cortex and their cell types using our single-cell transcriptomic atlas for PPC as well as independent data for different cortical regions^5^. In addition to PPC, our analysis identified genetic association between intelligence genes and multiple other cortical regions including ENT, a region involving in learning and memory. Interestingly, while there are intelligence genes expressed in multiple cortical regions, there are a significant number of the intelligence genes expressed specifically in a given cortical region. This raises the possibility that region-specific intelligence genes could contribute distinctively towards different abilities. Indeed, our data shows that mice with PPC or ENT-specific expressed intelligence genes display corresponding quantification or learning memory defects, but not both. Furthermore, the patients with PPC resection lesions in the left hemisphere display quantification defects, but not other cognitive abilities. Together, our study has, for the first time, provided experimental data supporting the concept of multiple intelligence abilities.

## MATERIALS AND METHODS

### PPC dissection

We used stereotaxic apparatus to pinpoint the PPC location for further tissue dissection. Isoflurane was used for anesthetization under the stereotaxic apparatus. Microinjector with needle fixed on the stereotaxic apparatus was used to set the location on the cranial bone surface after which, the cranial bone covering the targeted PPC region was ground off by mouse skull drill with a tip size of 0.5mm. When the surface of the PPC region was fully exposed, the mouse was euthanized by cervical dislocation. The exposed PPC region was cut and dissected by RWD mini scalpel (Cat# S33007-12) and quickly frozen in liquid nitrogen. Tissue was stored at −80°C for further single-nucleus extraction and RNA-sequencing. For each replicate, twenty-five C57/BL6J mice of seven weeks old were sacrificed for single-nucleus RNA-sequencing.

PPC location coordinates from bregma in millimeter with a rectangular area including two hemispheres: AP from −0.7 to −3.0 and ML from +3.2 to −3.2. This rectangular area occupies the central position (bregma: posterior 1.7-2.0 mm, lateral: 1.5-1.7mm) of PPC^26,27^.

### Single-nucleus RNA-sequencing

The cell suspension was loaded into Chromium microfluidic chips and barcoded with a 10× Chromium Controller. RNA was subsequently reverse-transcribed and sequencing library were constructed with reagents from a Chromium Single Cell reagent kit according to the manufacturer’s instructions (10× Genomics), targeting 10,000 cells per replicate. Sequencing was performed using Illumina NovaSeq to a depth of ∼80,000 reads per cell. Raw data were analyzed using Cell Ranger and Seurat^28^.

### Single-molecule Fluorescent in situ hybridization (smFISH)

Seven weeks old C57/BL6J mice were fully anesthetized by avertin. Anesthetized mice were firstly perfused by PBS transcardially to eliminate body blood, especially the blood in the brain and then perfused by 4% paraformaldehyde (PFA). The collected mouse brains were postfixed in 4% PFA for 6-8 hours at room temperature. Washed by PBS for three times, the brain tissues were embeded in OCT and quickly frozen in −80 °C storage freezer. The frozen samples were transferred to −20 °C to recover for 30 minutes and then coronally cut into 15 mm thick sections at −21 °C using a cryostat (Leica CM 1950). The brain sections containing PPC region were collected and stick to amino silane coated glass slides and stored at −80 °C for further processing. For FISH experiments, RNAscope® Multiplex Fluorescent Reagent Kit v2 was used (Advanced Cell Diagnostics, Cat# 323100). Briefly, frozen slides containing brain sections were immersed in precooled neutral buffered 10% formalin solution and incubated at 4 °C for 15 minutes. Then the samples were gradually dehydrated in 50%, 70% and 100% ethanol for five minutes respectively and dried for five minutes. Ten minutes permeabilization with H_2_O_2_, the samples were treated with Protease III. Then the probes were applied on the samples for hybridization followed by three steps of sequential amplification and application of fluorophores. DAPI (Sigma Cat# D9542) was used to stain the cell nucleus. Slides were mounted with Fluoroshield (Sigma Cat# F6182) and covered by coverslip for further imaging. RNAscope probes used include: *Npnt* (Advanced Cell Diagnostics, Cat# 316771), *Cux2* (Advanced Cell Diagnostics, Cat# 469551-C2), *Rorb* (Advanced Cell Diagnostics, Cat# 444271-C3), *Deptor* (Advanced Cell Diagnostics, Cat# 481561-C3), *Foxp2* (Advanced Cell Diagnostics, Cat# 428791-C2), RNAscope® 3-plex Positive Control Probe (Advanced Cell Diagnostics, Cat# 320881), RNAscope® 3-plex Negative Control Probe (Advanced Cell Diagnostics, Cat# 320871). Slides were scanned on an Akoya Vectra Polaris Scanner (Akoya Biosciences) using standard protocol provided by the instrument supplier, which generates a single unmixed whole slide scan containing multiple fluorescent channels. Images were further opened and processed by Phenochart 1.1.0 software.

### Human traits association analysis using MAGMA

We used MAGMA ^18^ to evaluate the association of a certain cell type with a particular human trait. A total of ten large-scale GWAS results of brain-related traits and disorders, including intelligence^20^, cognition^21^, memory^21^, Schizophrenia^11^, Parkinson’s disease^17^, Alzheimer^13^, attention deficit hyperactivity disorder (ADHD)^12^, bipolar disorder^14^, depression^15^ and obsessive-compulsive disorder (OCD)^16^ were included.

Sporadic or very low gene expression will lead to a false discovery of highly specific cell marker. We thus excluded low-expressed genes with total expression levels below 0.01 in all cells. MAGMA_Celltyping (https://github.com/neurogenomics/MAGMA_Celltyping) was then used to calculate cell-type specificity and detect a positive association (one-sided test) between the gene-level traits associations and the cell-type specificity.

### Food-based quantification assay of mice

Mice were placed at the center of a 30×30×50cm open field arena at the start of the test and allowed to move freely for about 10 min for environment adaption. The food was then placed into the two containers located in opposite corners of the arena (20g:80g). Individual mouse movements were captured using a video camera over a period of 60 min. Wild type mice prefer to choose the corner with more (80g) food. The ratio of accumulated duration in the zones with more (80g) and less (20g) food was calculated to assess quantification ability.

Seven weeks homozygous *Rnf123^-/-^* (GemPharmatech Co., Ltd# T045030), *Atp2a1^+/-^* (GemPharmatech Co., Ltd#T012468) and wild-type mice (C57BL/6J) were used to perform food-based quantification assay. Individual mice were restrained from food for 48 hours before experiments.

### Mouse Y maze test for working memory

Seven weeks homozygous *Rnf123^-/-^*, heterozygous *Atp2a1^+/-^,* and wild-type mice were used to evaluate their working memory by Y-maze test. Animals were brought from home cage to the test room and allowed for habituation for 1h. The test was performed in a Y-shaped maze with three 30cm dark-colored arms orientated at 120° from each other. Individual mouse was placed gently at a fixed position in the maze and allowed for freely exploring for 5 minutes. Distance moved, moving speed and arm entries were recorded by Ethovision XT (Noldus, the Netherlands). An alteration was defined as consecutive entries in all three arms. Working memory was assessed by spontaneous alterations as previously described^29^.

### Morris Water Maze test

Seven weeks mice were firstly tested for cued learning 1 day prior to the test, where the hidden platform was slightly above the water surface with a cued black object mounted. Mice were allowed to locate the escape platform with the visible cue and successfully climb on within 1 minute. Basic abilities such as sensorimotor ability, intact eyesight, and motivation were evaluated with escape latency and swimming velocity.

Mice were then trained to find a 10cm diameter circular hidden platform submerged 1 cm below the surface in a 120cm diameter circular tank filled with white milk water at 22°C. Four black A4-size shapes around the tank served as visual cues (hexagon, circle, triangle, rectangle). Mice were trained with 4 trials per day with a 30-minute interval for 5 days. Among the 4 trials, mice were gently released from 4 different locations facing the wall and were allowed to freely explore for 1 minute. A trial ended when mice reached the platform and stayed for 1 second, after which mice remained on the platform for 15 seconds to facilitate learning. Unsuccessful mice were guided to the platform and left for 15 seconds. Mice that failed to reach the platform were recorded with a 1-minute escape latency. The first trial on the first day was recorded as day 0. Learning and memory ability were assessed by average escape latency to the platform.

All the data were recorded by tracking software EthoVision XT 15 and analyzed by Graphpad Prism 8.0.

### Human logic, language and working memory tests

We used three tests as previous study^20^, including reasoning, language and working memory test. Each test contains 10 questions. All questions were derived from UK Biobank Fluid Intelligence Test^6^, Raven’s Progressive Matrices^30^, STR-MADT (https://www.sdu.dk/en) and the Mainland Chinese Version of the Wechsler Adult Intelligence Scale (WAIS-RC)^31^. The scores represented the number of correct answers out of 10 questions.

## Acknowledgements

This research was supported in part by the grants from National Natural Science Foundation of China (U21A20201) and the Science Technology Department of Zhejiang Province (2020E10027, 2021ZY1019, 2022ZY1005), the grant for the construction of Key Laboratory of Growth Regulation and Translational Research of Zhejiang Province, Zhejiang Leading Innovative and Entrepreneur Team Introduction Program (2018R01003) and Research Program No. 202208011 of Westlake Laboratory of Life Sciences and Biomedicine.

## Author contributions

JJ.L. and R.C. contributed to sample preparation for snRNA-seq and FISH experiment.

D.Y. conceived the methods for computational analysis. D.Y. and X.Y. preformed bioinformatic analysis. J.P. and L.X. performed mouse behavioral assay and characterized quantification ability. S.G. performed mouse Y maze and Morris Water Maze tests. J.P., S.G., L.X., C.X., X.H. and JF.L. completed human tests. T.X. and D.Y. wrote the paper. T.X. and D.Y. designed and advised on the whole project.

## Competing Interest Declaration

Authors declare that they have no competing interests.

---

**Figure S1.**
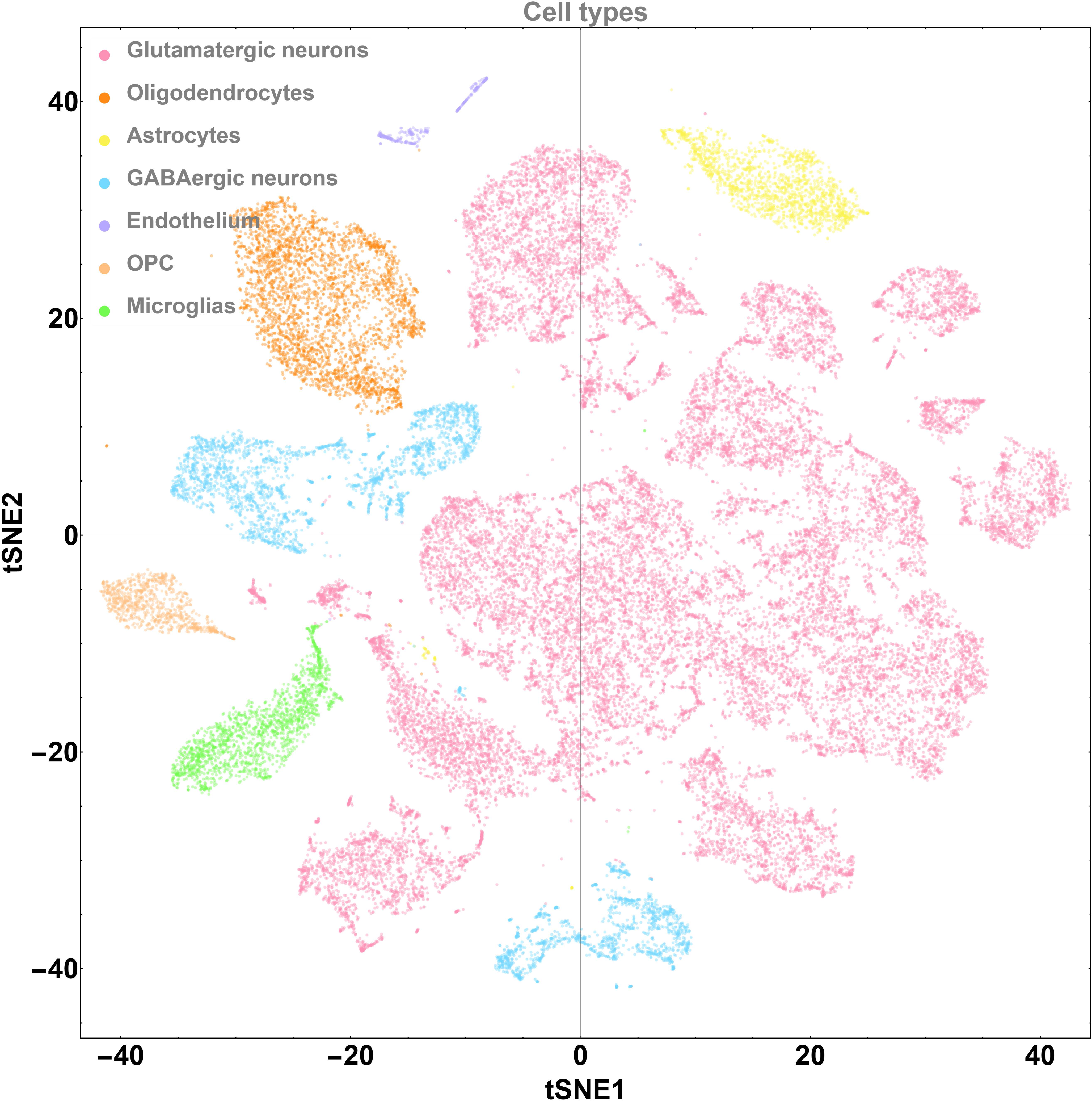
Profiling of PPC cell types. t-SNE plot visualization of the PPC cell types. Dots represent cells, and distances between dots reflect transcriptomic similarity. Cell are colored by morphological types.

**Figure S2.**
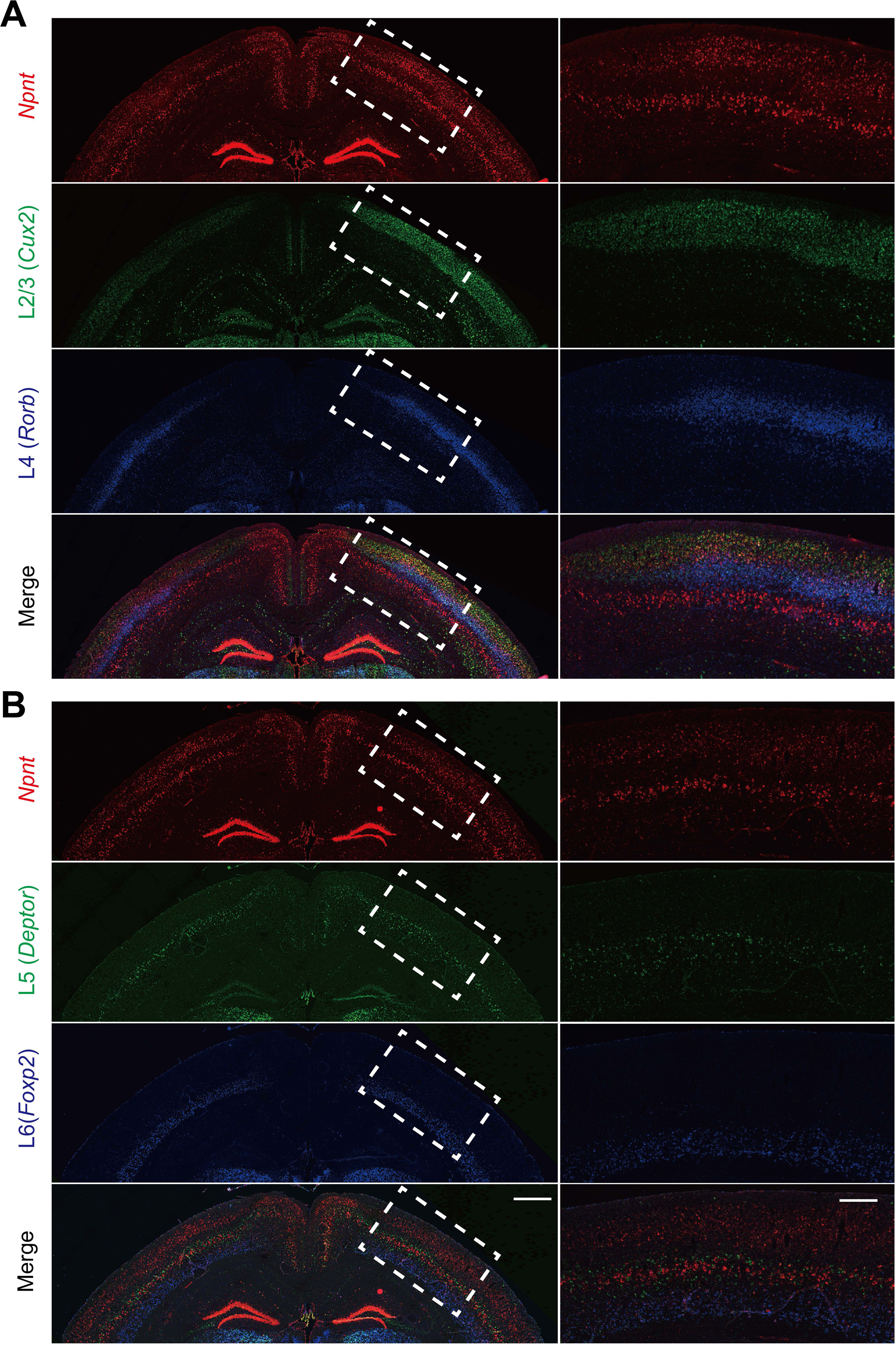
**Validation of *Npnt* neurons**. FISH images showing *Npnt* neurons located in the layer 2/3 (A) and layer 5 (B). *Npnt* expressing neurons are different from *Cux2* expressing L2/3 or *Deptor* expressing L5 excitatory neurons. Scale bars: left, 800 μm; right, 300 μm.

**Figure S3.**
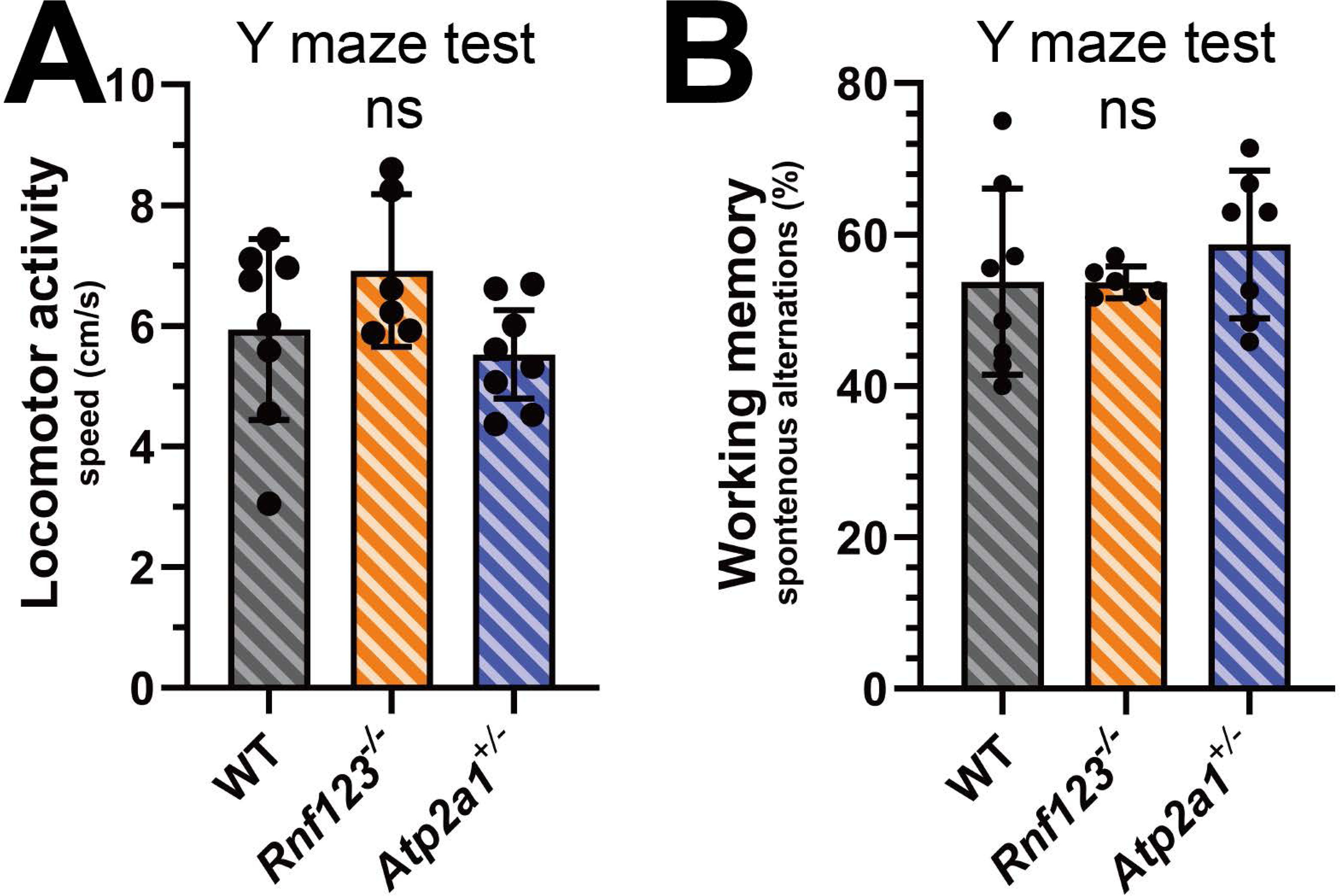
Analysis of the mouse behaviors in y maze. *Rnf123* and *Atp2a1* mutants show normal motor function (**A**) and working memory (**B**).

**Figure S4.**
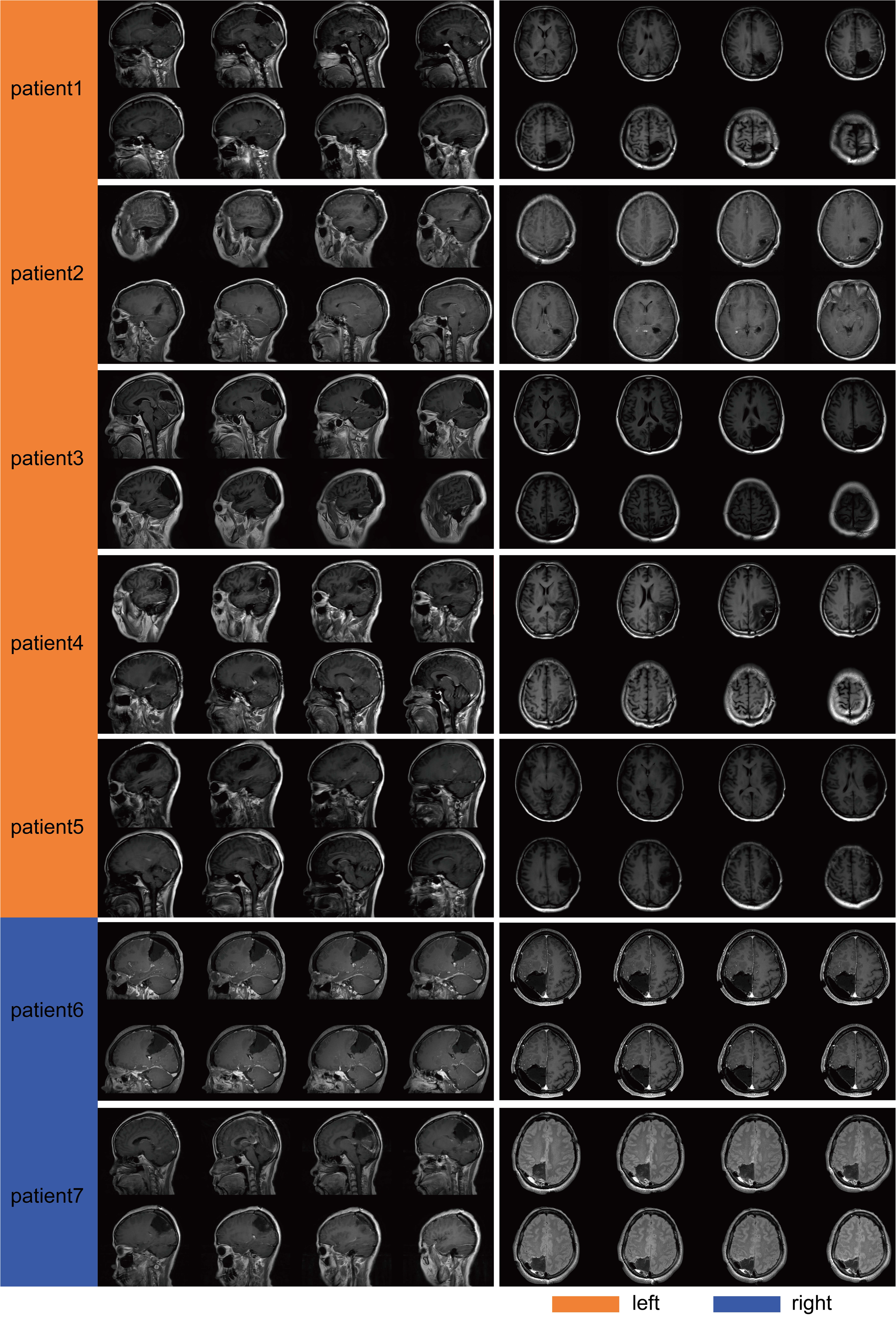
**Sagittal and axial magnetic resonance imaging scan showing brain resection of 7 participants.**

## Notes

### Competing Interest Statement

The authors have declared no competing interest.

